# Parental effects in a filamentous fungus: phenotype, fitness, and mechanism

**DOI:** 10.1101/2022.12.09.519717

**Authors:** Mariana Villalba de la Peña, Pauliina A. M. Summanen, Neda N. Moghadam, Ilkka Kronholm

## Abstract

Adaptation to changing environments often requires meaningful phenotypic modifications to match the current conditions. However, obtaining information about the surroundings during an organism’s own lifetime may only permit accommodating relatively late developmental modifications. Therefore, it may be advantageous to rely on inter-generational or trans-generational cues that provide information about the environment as early as possible to allow development along an optimal trajectory. Transfer of information or resources across generations, known as parental effects, is well documented in animals and plants but not in other eukaryotes, such as fungi. Understanding parental effects and their evolutionary consequences in fungi is of vital importance as they perform crucial ecosystem functions. In this study, we investigated whether parental effects are present in the filamentous fungus *Neurospora crassa*, how long do they last, are the effects adaptive, and what is their mechanism. We performed a fully factorial match / mismatch experiment for a good and poor quality environment, in which we measured mycelium size of strains that experienced either a matched or mismatched environment in their previous generation. We found a strong silver spoon effect in initial mycelium growth, which lasted for one generation, and increased fitness during competition experiments. By using deletion mutants that lacked key genes in epigenetic processes, we show that epigenetic mechanisms are not involved in this effect. Instead, we show that spore glycogen content, glucose availability and a radical transcription shift in spores are the main mechanisms behind this parental effect.

## Introduction

Organisms are obligated to adjust their phenotype throughout development to match the current conditions. However, phenotypic changes triggered by the environment during an organisms own lifetime might only permit to accommodate relatively late developmental modifications. Therefore, it may be beneficial for an organism to obtain cues or resources from the parents, as both generations are likely to face similar environmental conditions. The effect that the parental phenotype, or environment, has on the offspring fitness is known as a parental effect [1].

Parental effects are usually studied using match / mismatch experiments. These are fully factorial reciprocal transplant experiments where the offspring performance is measured in the same or different conditions compared to their parental environment [2]. This experimental design is necessary to determine the existence and the type of parental effects. One possible outcome is adaptive matching, or anticipatory effects, meaning that offspring performance is greater when its own environment matches the parental environment. Another possible outcome is a carry over, or silver spoon effect. This happens when the quality of the parents, or the parental environment, is the main factor that shapes offspring fitness. Parental effects can also be a combination of adaptive matching and carry over effects [2].

There are many examples of parental effects in plants and animals. For example, in the crustacean *Daphnia* a signal perceived by the mother that induces the development of a defensive structure can be inherited from mothers to offspring [3]. In some plants, the light environment of the mother influences the fitness of offspring [4], in the plant *Arabidopsis* offspring can inherit responses to osmotic stress [5], or pathogens [6]. Furthermore, it is increasingly suggested that parental effects may contribute to adaptive evolution [1, 7, 8, 9].

Parental effects can be transmitted via several mechanisms. One of them is the quality and quantity of provisional molecules such as nutrient reserves, mRNAs, and proteins. These supplies could be altered by the parental condition and significantly impact the offspring’s performance either at early, or all stages along its lifetime [10, 11, 12]. Also, parental conditions can induce epigenetic changes (e.g DNA methylation and histone modifications) which can be inherited in some cases and influence gene expression and phenotypic traits [5, 10, 13, 14, 15]. For instance, in dandelions DNA methylation patterns induced by environmental stressors can be transmitted to the next generation even when the stressor is removed from the offspring environment [16]. In plants, the RNA directed DNA methylation pathway has been implicated in inherited parental effects [5, 17]. The mechanism behind the parental effect may influence its duration. If the underlying mechanism is epigenetic, the parental effect may persist across generations, while a provisioning effect may be only brief [10]. Even though parental effects have been widely studied, the underlying mechanisms are rarely documented [18]. To understand how parental effects aid adaptation it is crucial to first understand under which circumstances parental effects manifest, their duration, and their underlying mechanisms.

So far, most of the research on parental effects has focused on plants and animals, and to date just a few a studies have investigated the existence of parental effects in microbes. Even though theoretical models suggest that parental effects are expected to evolve when environmental fluctuations span several generations, which may be often true for microbes [19]. To our knowledge only one previous study has investigated maternal effects in a fungus. Zimmerman et al. in 2016 reported the existence of asymmetrical investment in *Neurospora crassa*. The authors discovered that, when the fungus reproduces sexually, maternal effects influences spore number and germination success [20]. The lack of research of parental effects on fungi is surprising, as they perform key ecosystem functions such as organic matter decomposition and are involved in plant symbiosis [21].

To understand parental effects in fungi, we investigated the existence and mechanisms of parental effects in the filamentous fungus *Neurospora crassa. N. crassa* has a facultative sexual cycle, but we focused on parental effects that are transmitted through asexual spores, called conidia. We also determined the fitness relevance and duration of such effects. Finally, we investigated the mechanisms behind the parental effects by quantifying nutrient reserves, using mutants, and RNA-seq. Our study is one of the first to thoroughly examine parental effects in fungi.

## Methods

### Existence of parental effects

To investigate whether parental effects exist in *N. crassa* we performed a reciprocal match / mismatch experiment [2]. We compared the initial mycelium growth in two different environments where the strains had experienced either the same or a different environment in the previous generation (Fig 1A). We inoculated conidia of *N. crassa* strain 2489, obtained from Fungal Genetics Stock Center [22], in slants containing Vogel’s medium N (VM) [23] with either 1.5% or 0.015% sucrose. The fungus grew in the slants for one generation, defined here as growing from spore to spore. Each slant represented a biological replicate. At the end of generation one, conidia were harvested and filtered to remove any mycelial fragments, then counted and measured using a CASY cell counter with a 45 µm capillary and a gating window of 2.5–10 µm. We inoculated 10 000 conidia at the center of a petri dish with VM agar, containing the parental or a different sucrose concentration. Plates were randomized and incubated at 25 ^*◦*^C. We measured the diameter of the colony at three time points: first after 14 to 18 hours from inoculation, then second and third measurements 2 to 4 hours apart from the previous measuring time. Growth rate was estimated as the slope of the linear regression of time against colony diameter. To make sure that differences in mycelial growth was not driven by spore viability or dormancy, we measured conidial viability by plating the harvested conidia on sorbose medium. Sorbose induces colonial morphology in *N. crassa* [24], this allowed us to count the number of germinated conidia after three days of incubation at room temperature. The experiment was repeated nine times, sample sizes for each experiment are reported in supplementary table S1.

**Figure 1:**
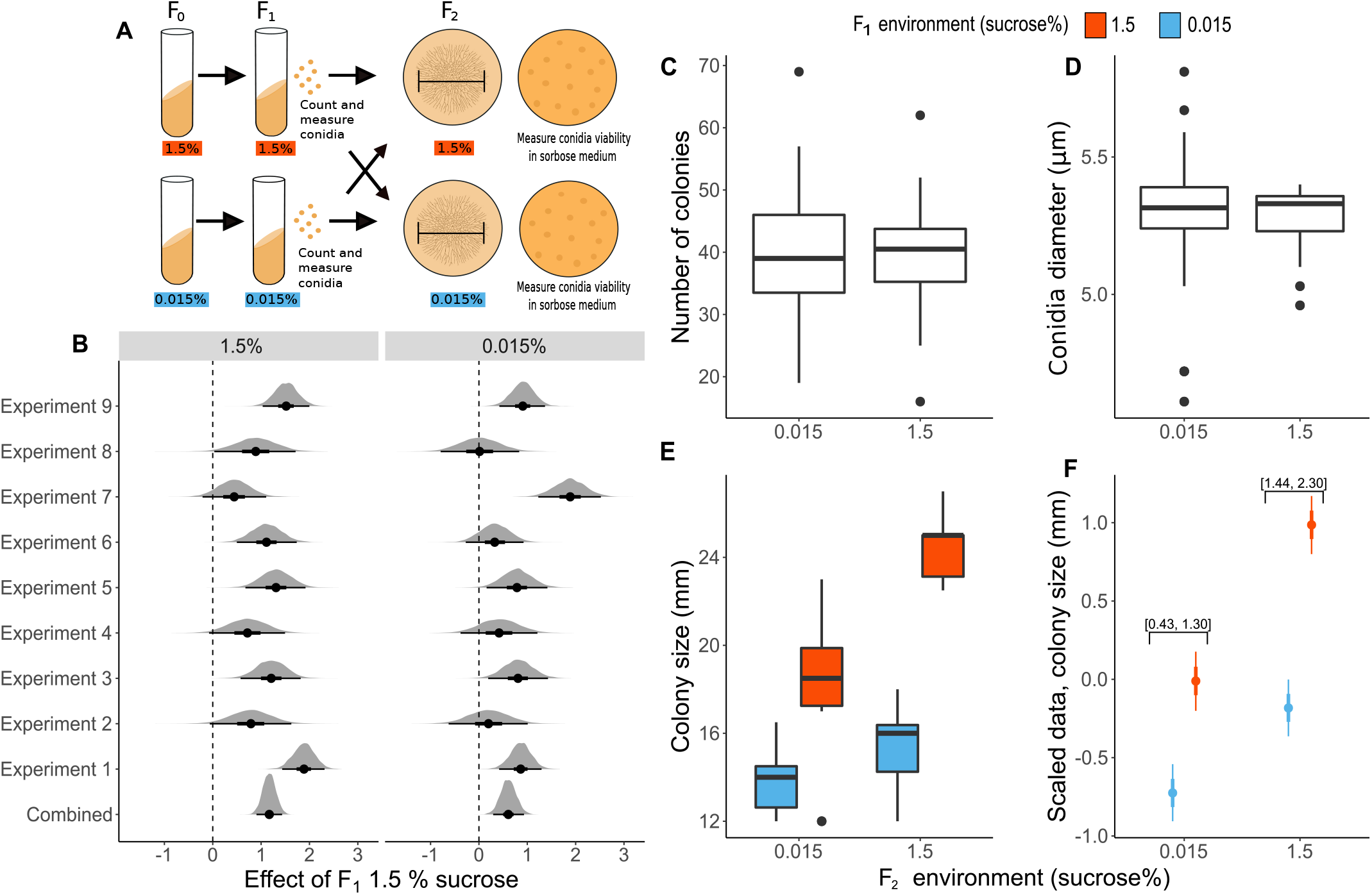
Effect of F_1_ sucrose concentration on colony size in F_2_. **(A)** Experimental design. Fungus was cultivated in 1.5% or 0.015% sucrose slants for two generations, then the same number of conidia were plated on plates with 1.5% or 0.015% sucrose. The mycelial diameter was measured and the number of colonies formed in sorbose plates was counted to estimate spore viability. **(B)** Posterior distributions of the effect of F_1_ 1.5% sucrose on initial colony size, when F_2_ was grown in either 1.5% or 0.015% sucrose. **(C)** Number of colonies in sorbose plates. **(D)** Conidial diameter. **(E)** Raw data of initial colony size from experiment one. **(F)** Model estimates using the combined data. The 95% HPDI of the difference between treatments is shown in square brackets.

We also explored the existence of parental effects on alternative carbon sources. We performed a match / mismatch experiment where we compared sucrose to an alternative carbon source: cellulose, lactose, maltose, or xylose. We measured initial colony size when the fungus was exposed to either the same or a different carbon source in the previous generation. There were five biological replicates for each treatment.

### Duration of parental effects

To investigate whether the parental effects persisted for more than one generation we continued the experiment described above, into the third generation. At the end of the second generation, conidia were harvested, counted and plated. Mycelial growth was measured in plates that either matched or mismatched the F_1_ sucrose environment (Fig 2A). We assessed conidia viability as above. We repeated this experiment five times, the sample size of each experiment is reported in supplementary table S1.

**Figure 2:**
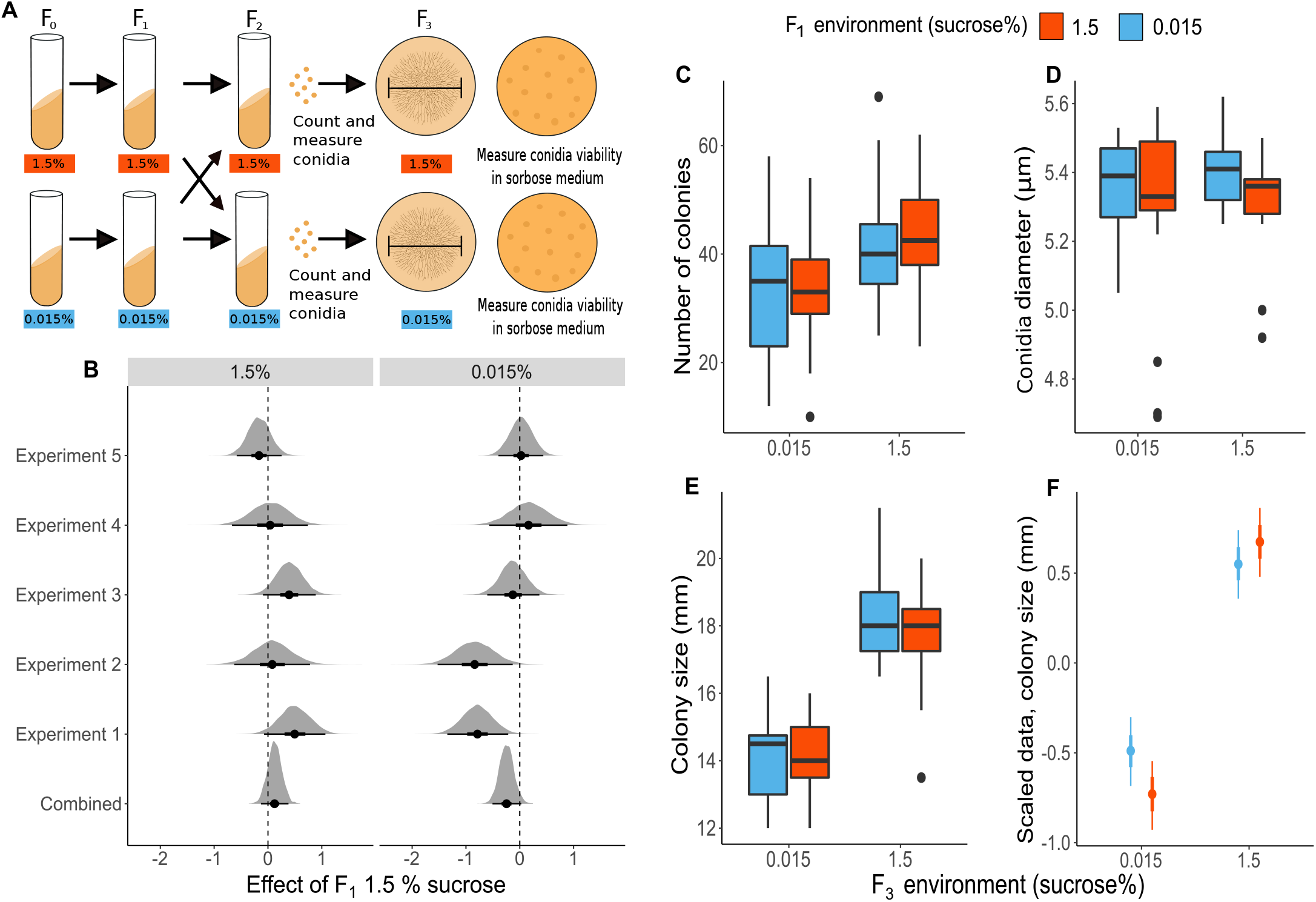
Effect of F_1_ sucrose concentration on colony size in F_3_. **(A)** Experimental design. Fungus was cultivated in 1.5% or 0.015% sucrose slants for two generations, then matched or mismatched for one generation, and then the same number of conidia were plated on plates with 1.5% or 0.015% sucrose. The mycelial diameter was measured and the number of colonies formed in sorbose plates was counted to estimate spore viability. **(B)** Posterior distributions of the effect of F_1_ 1.5% sucrose on initial colony size, when F_3_ was grown in either 1.5% or 0.015% sucrose. **(C)** Number of colonies in sorbose plates. **(D)** Conidial diameter. **(E)** Raw data of initial colony size from experiment one. **(F)** Model estimates using the combined data.

### Fitness consequences of parental effects

To estimate the fitness effects that the parental environments cause, we used a competition experiment with marked strains. We have previously developed marked strains for *N. crassa* by inserting a DNA barcode to the *csr*-locus, this marker allows us to estimate the proportion of the marked strain in a sample of conidia using high resolution melting (HRM) PCR [25]. The experimental design was the same as in the initial match / mismatch experiment, except instead of plating conidia at the F_2_ generation, we combined conidia from two different strains in a slant, and let them produce conidia in competition. Then we estimated the proportion of the marked strain in the conidial sample using HRM-PCR (Fig S4A). We have previously observed that the mating type locus and the *csr*-tag have fitness effects, so in order to estimate the fitness effect of the parental sucrose environment we combined the parental environment, competition environment, the *csr*-tag, and the mating type locus in 8 different combinations (Table S4). Strains with the same mating type were never competed against each other, because in these nearly isogenic strains, hyphae of the same mating type would fuse together and no competition would occur [25]. A detailed description of the competition experiments can be found in supplementary methods.

### Mechanisms of parental effects

#### Protein content and carbohydrate reserves

To investigate qualitative differences in the conidia, we assayed whether protein, glycogen, or glucose reserves differed between F_2_ conidia coming from 0.015% or 1.5% sucrose. We measured protein and sugars using kits: BCA protein assay kit (Thermo Scientific), glycogen assay kit (Sigma-Aldrich, MAK016), and glucose assay kit (Sigma-Aldrich, MAK263), accoring to manufacturers’ instructions. We extracted total protein from 40 million conida, and glycogen and glucose from 70 million conidia (see supplementary methods).

#### RNA-seq of conidia

To understand the mechanisms behind parental effects we investigated the gene expression patterns of F_2_ conidia cultured in either 1.5 or 0.015% sucrose. We extracted RNA from conidia following [26]. See supplementary methods for details and supplementary table S3 for purity, concentration and integrity metrics of the extracted RNA. We used the ERCC RNA Spike-ins [27] as external controls (see supplementary methods). Six biological replicates from each sucrose concentration were sent to Novogene for mRNA poly A enrichment library preparation and transcriptome sequencing, using the Illumina NovaSeq platform with 150 bp paired-end libraries.

#### RNA-seq normalization and analysis

We examined the quality control metrics of the twelve sequenced samples with FastQC. The samples were aligned against *N. crassa* reference genome (assembly NC12) with the added 92 ERCC RNA Spike-In control transcripts. We aligned the sequences using hisat [28] specifying 2 500 as the maximum intron length [29], all other parameters were set as default. The number of obtained reads and alignment metrics are reported in the supplementary table S3.

We normalized the data using the Trimmed mean of the M-values approach, and then used the ERCC spike in controls to remove unwanted variation using the RUVseq package [30]. Finally we used DESEq2 to obtain differentially expressed genes and cluster profiles to perform an over representation analysis (ORA) and a gene set enrichment analysis (GSEA). See supplementary methods for details.

#### Epigenetic mechanisms

To investigate whether parental effects relied on epigenetic mechanisms we performed the same basic match / mismatch experimental design (Fig 1A), but with three deletion mutants deficient for different epigenetic mechanisms. The mutants were: ∆*dim-2* which lacks DNA methylation [31], ∆*qde-2* which has compromised RNA interference pathway [32], and ∆*set-7* which lacks trimethylation of the lysine 27 on the histone 3 (H3K27me3) [33]. Sample size was *n* = 40 for each mutant strain. The mutant strains have been previously described in Kronholm et al. 2016 [34].

We further explored the overlap between the genes belonging to the main GSEA enriched pathways and different genomic domains. In *N. crassa* trimethylation of histone 3 lysine 9 (H3K9me3) is associated with heterochromatin, H3K27me3 with facultative heterochromatin, and dimethylation of histone 3 lysine 36 (H3K36me2) with euchromatin. DNA methylation occurs only in H3K9me3 domains. Briefly, we obtained ChIP-seq reads for each of the domains, we align them to the reference genome and identified the domains of the histone modifications (see supplementary methods).Then we identified the intersecting regions between each histone modification domains and the genes from each GSEA enriched pathway. We considered a gene to belong to a histone modification domain if at least 20% of the gene overlapped with the histone modification domain.

### Statistical analyses

#### Existence and duration of parental effects

Since time of the first measurement varied between experiments, the data was centered and variance standardized experiment by experiment. We fitted Bayesian models using Hamiltonian Monte Carlo implemented with the Stan language [35] using the “ulam” function available in the rethinking package [36], in R version 4.0.2. We fitted a model with initial colony size as response, treatment and spore viability as predictors, and the slant as a random factor. See supplementary material for details. The estimates and the highest posterior density intervals (HPDI) of all models are reported in the supplementary table S5.

For both F_2_ and F_3_ data, we analyzed each experiment separately and for all experiments combined. (Figure 1B & 2B). When analyzing each experiment independently we did not considered slant (*β*_*s*_) an viability (*β*_*c*_) in the model because the sample was not big enough for the model to converge. When analyzing initial growth of F_2_ we did not considered viability (*β*_*c*_) in the model as it did not have a significant effect and three experiments had missing viability data.

#### Fitness consequences of parental effects

We estimated the relative fitness effect of the parental 1.5% sucrose environment following the same principle as in Kronholm et al. 2020. We used a model that takes uncertainty in proportion estimates of the marked strain into account, and models the log-ratio of the strain proportions. With this model specification the slope of the model is log of relative fitness [25]. The log-ratio of the strain proportions was the the response, effect of *csr-1\** marker, mating type, and parental environment were predictors, and population was a random factor. See supplementary material for details.

## Results

### Existence of parental effects

To test if different parental resources cause parental effects in *N. crassa*, we performed a reciprocal match / mismatch experiment with a rich (1.5%) and a poor (0.015%) sucrose environment. All the results from the Bayesian model in equation S1 are reported as means with 95% highest posterior density intervals (HPDI) in square brackets. We observed that initial size of the mycelial colony was always higher in the 1.5% sucrose environment (Figure 1E & F). Also, if the fungus experienced 1.5% sucrose in the previous generation, the initial size was higher regardless of the F_2_ assay environment. If the fungus experienced 1.5% sucrose in the previous generation, the initial colony size was 17% bigger when growing in 1.5% and 10% bigger when growing in 0.015% sucrose, both compared to the fungus growing in the same sucrose concentration but which experienced 0.015% sucrose in the previous generation. The difference for F_1_ treatments was 1.167 [0.913, 1.425] when grown in 1.5% sucrose, and 0.714 [0.450, 0.970] when grown in 0.015% sucrose. Since the parental environment with 1.5% sucrose always produces larger colonies in the next generation no matter what the current environment is, the parental effects observed here are due to 1.5% sucrose just being a better environment overall, with no evidence of any adaptive response to low resources by the fungus. This type of parental effect is also called a silver-spoon effect, since an individual growing in a better environment will always be better off [37].

We repeated the experiment nine times. We observed some variation in experimental outcomes for unknown reasons. In some of the experiments, the effect of the F_1_ environment overlapped with zero but when data from all experiments was combined and analyzed together there was a clear effect of the parental environment (Fig 1B, 1F).

*N. crassa* produces around 11 times less conidia when sucrose concentration is 0.015% (Figure S1), the difference for scaled data was 1.743 [1.519, 1.974]. It is known that number of germinating conidia affects the rate at which the mycelium develops [38]. Therefore, we always counted conidia and plated the same number of conidia on plates. To make sure that differences in conidial viability or dormancy induced by the different treatments were not a factor, we measured the number of colony forming units in our samples by plating. We did not observe any differences in conidial viability in any generation for conidia coming from either 1.5% or 0.015% sucrose, difference in F_2_ was *−*0.174 [*−*0.712, 0.379], and in F_3_: 0.209 [*−*0.808, 0.419] (Fig 1C and 2C). Therefore, there must be some qualitative difference in the conidia originating from 1.5% and 0.015% sucrose. We checked if spore size was different, but we did not observe any differences: difference in size in F_2_ samples was *−*0.019 [*−*0.372, 0.305], and for F_3_ samples *−*0.423 [*−*1.103, 0.225] (Fig 1D, 2D).

We also screened alternative carbon sources for the presence of parental effects. We performed the same experiment but compared the 1.5% sucrose environment against 1.5% arabinose, cellullose, lactose, maltose or xylose. We found a similar silver spoon effect when *Neurospora* was grown with arabinose, cellulose or lactose. Difference in initial colony size when the strain was grown in sucrose versus arabinose in F_1_ was 1.841 [1.198, 2.462]; for cellulose difference was 1.496 [0.973, 2.056]; and for lactose 1.809 [1.159, 2.451] (Fig S3). In each of these environments we observed that fungus grew always bigger when it experienced sucrose during the previous generation. When comparing sucrose to maltose or xylose we did not observe any parental effects (Fig S3).

#### Duration of parental effects

Next, we estimated the duration of the observed parental effect by continuing the experiment to F_3_ (Fig 2A). The silver spoon effect observed in F_2_ did not carry on to subsequent generations. The F_1_ environment did not have an effect on initial growth in F_3_, the effect of F_1_ environment in 1.5% sucrose was 0.121 [*−*0.122, 0.390], and *−*0.240 [*−*0.499, 0.011] in 0.015% sucrose. We repeated the experiment five times, as in the F_2_ experiment we observed some variation in experimental outcomes. In some of the experiments, there appears to a significant F_1_ effect in cultures with 0.015% sucrose. However, when all experiments were combined the effect of the F_1_ environment overlapped with zero (Fig 2B, 2F).

We further investigated the duration of the silver spoon effect by looking the growth rate of the mycelium on F_2_ plates in more detail. We had taken three measurements of the colony size on the F_2_ plates. When we calculated growth rates instead of using initial colony size, we observed that F_1_ environment only had an effect on the growth rate calculated from first time points, and no effect on growth rate in the subsequent time points (Figure S2). Taken together, these experiments suggest that the observed parental effect is an intergenerational effect that matters in the establishment of the mycelium. As the mycelium grows in size, the effect disappears.

### Fitness consequences of parental effects

Next, we wanted to understand the biological significance of the observed silver spoon effect, by investigating how does parental environment contribute to offspring fitness. We performed the match / mismatch experiment as before, but instead of plating the conidia we combined the conidia from two strains and let them compete (Fig S4A). We found that the relative fitness of a strain that experienced 1.5% sucrose environment in the previous generation was approximately four times higher when competing against a strain that experienced 0.015% sucrose in the previous generation, in both 1.5% and 0.015% sucrose competition environments (Table 1). This suggests that the small increase in initial speed of colony establishment matters greatly for fitness.

**Table 1:** Relative fitness effects estimated from competition experiments. Values below 1 indicate that fitness is decreased relative to the other genotype or parental environment, while values above 1 indicate higher relative fitness. The fitness effect of *csr-1\** is relative to wild type allele, *mat A* relative to *mat a* and F_1_ 1.5% is relative to 0.015% sucrose in the parental environment.

| Effect | $W_{ij}$ [95% HPDI] | | |
| --- | --- | --- | --- |
|  | Combined | F<sub>2</sub> 1.5% sucrose | F<sub>2</sub> 0.015% sucrose |
| *csr-1* * | 0.58 [0.5, 0.67] | 0.51 [0.4, 0.63] | 0.66 [0.55, 0.81] |
| *mat A* | 0.83 [0.72, 0.94] | 0.86 [0.7, 1.06] | 0.79 [0.65, 0.96] |
| F<sub>1</sub> 1.5% sucrose | 4.13 [3.41, 5.14] | 4.72 [3.52, 6.52] | 3.62 [2.82, 4.88] |

### Mechanisms of parental effects

Next, we explored possible mechanisms for the observed parental effects. We investigated nutrient composition, mRNA content of condia, and possible epigenetic effects.

#### Protein content and carbohydrate reserves

We quantified protein, glycogen and glucose content in the F_2_ conidia grown in either 1.5% or 0.015% sucrose. We observed no difference in the total protein content between treatments, scaled difference was *−*0.187 [*−*0.701, 0.311] (Fig 3A). However, we found that spores originating from 1.5% sucrose had a higher amount of glycogen, scaled difference was 1.59 [1.009, 2.206]; and a higher amount of glucose, scaled difference was 1.794 [1.44, 2.139] (Fig 3B, 3C). This suggest that carbohydrate storage in conidia may be responsible for the silver spoon effect.

**Figure 3:**
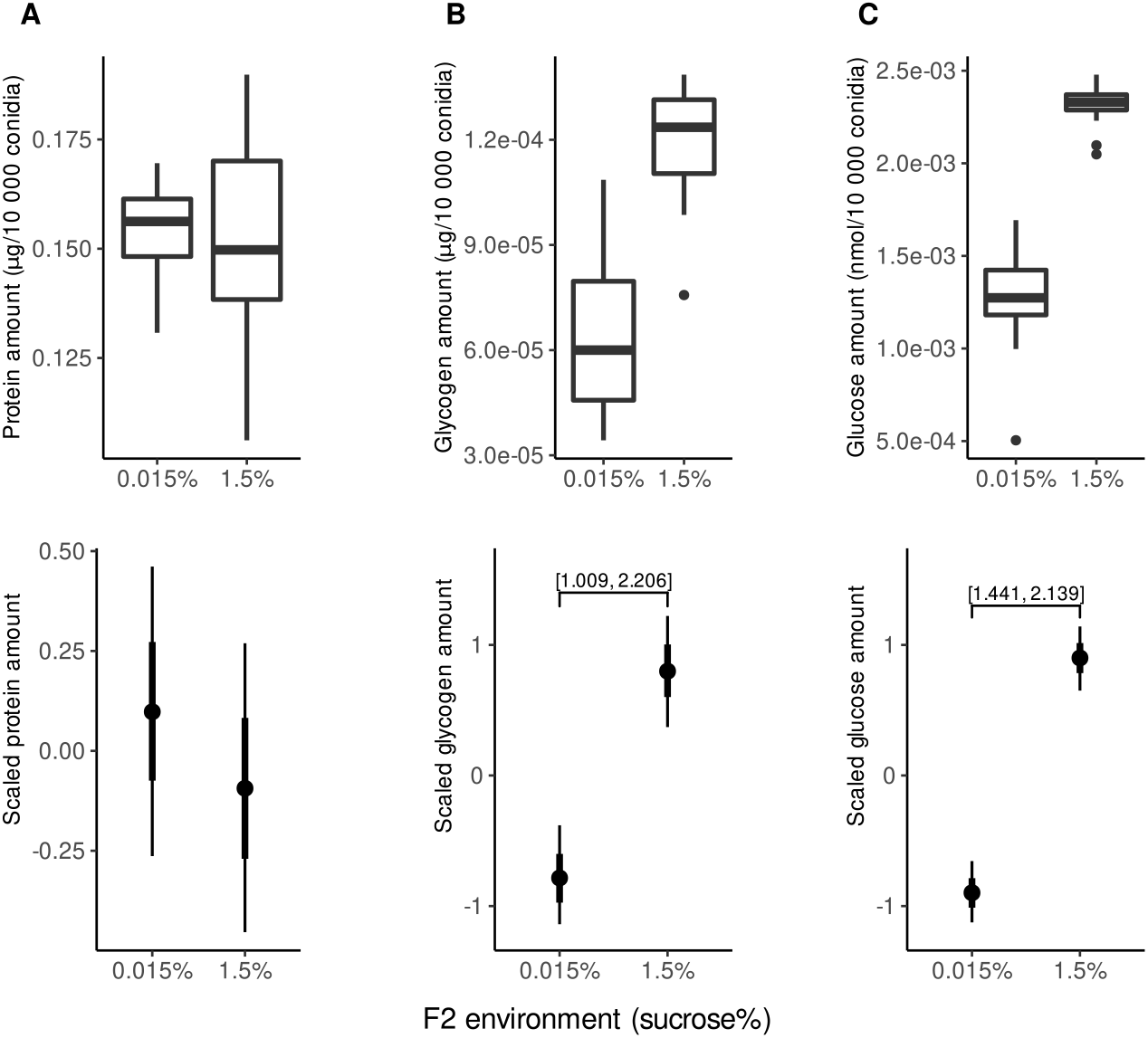
Effect of F_1_ environment on total amount of protein, glycogen and glucose in conidia. Raw data (top) and the model estimates (bottom) of the total amount of **(A)** protein **(B)** glycogen and **(C)** glucose in conidia originating from 1.5% or 0.015% sucrose. The data in the bottom row is scaled, numbers inside square brackets show the 95% HPDI of the difference between treatments.

#### RNA-seq of conidia

To further understand the physiological changes in conidia originating from 1.5% or 0.015% sucrose, we sequenced conidial mRNAs. On average we obtained 13 *×* 10^6^, of 150 bp reads per library (Table S3). More than 93% of reads in all the samples successfully mapped the reference genome (Table S3). Grouping samples by PCA showed that PC1 represented variation between the sucrose environments, and explained 63.86% of the variation, while PC2 represented variation across samples of the same treatment, and explained 12.37% of the variation (Fig 4A). We also observed a symmetrical distribution of differential gene expression where 6564 (p-adjusted < 0.01) of the 8925 annotated genes were differentially expressed between treatments (Fig 4B). The p-value distribution obtained from DESEq2 analysis is shown in figure S5D.

**Figure 4:**
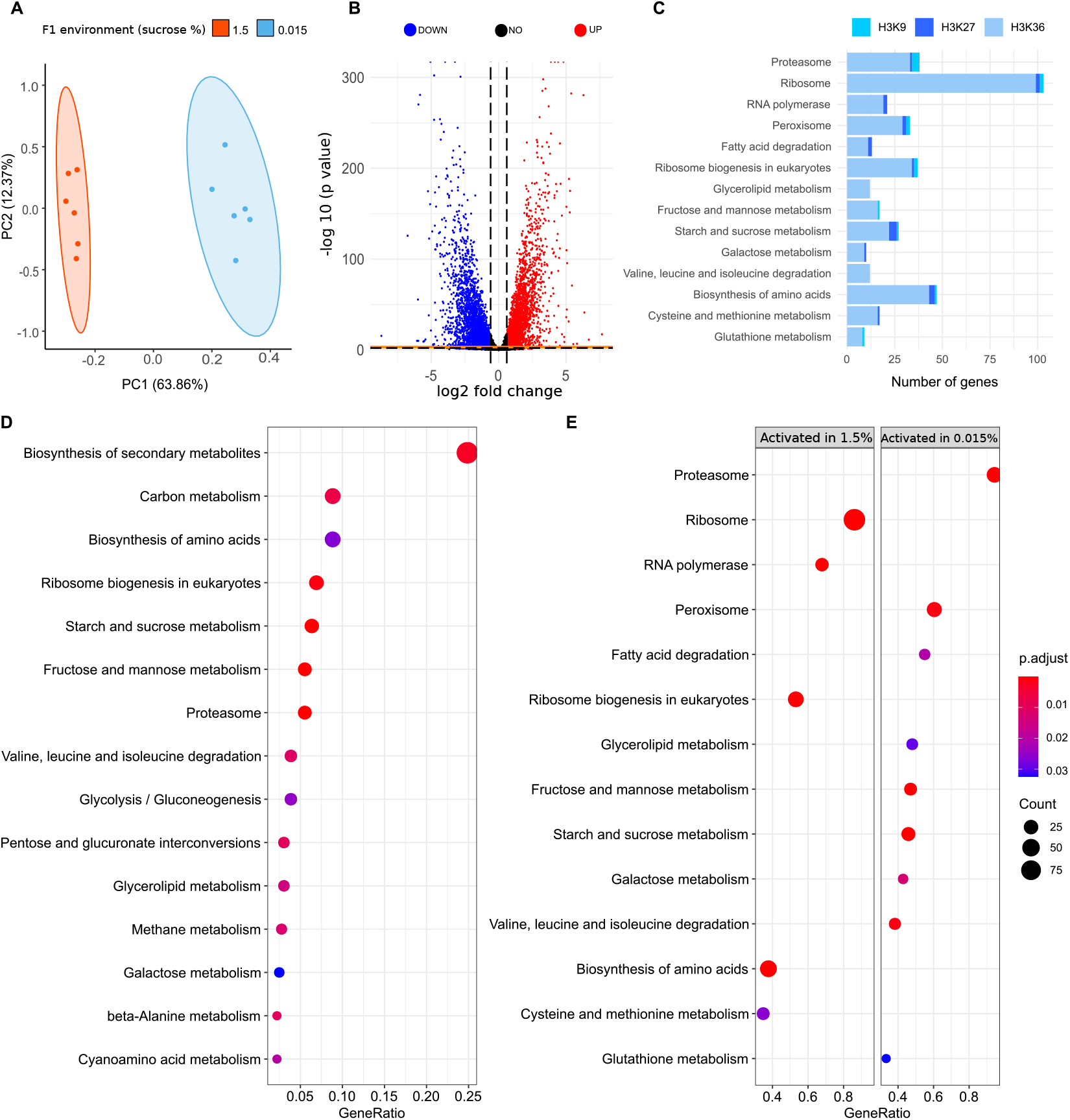
DE and enrichment results. **(A)** Principal component analysis of the read count data. **(B)** Volcano plot shows the distribution of the DE genes between treatments. Blue dots are downregulated genes, red dots are upregulated genes, and black dots are genes that were not differentially expressed. The horizontal dashed line and the solid orange line indicates the p-value of 0.01 and 0.001 respectively after correcting for multiple testing, the vertical dashed lines represent log fold changes of 1.5. **(C)** Number of genes of the most enriched pathways associated to three different histone modification domains: H3K9me3, H3K27me3 and H3K36me2. **(D)** KEGG enrichment pathways from over representation analysis (ORA). **(E)**. KEGG enrichment pathways results with gene set enrichment analysis (GSEA). The color gradient shows the p-value and dot size the count of genes in each pathway.

We performed two different enrichment analyses: an over representation of analysis of KEGG pathways, and a gene set enrichment analysis. Even though the two types of enrichment analysis show slight differences, all of the enriched pathways fall into three categories: metabolism, genetic information and processing, and cellular processes (Fig 4 D & E). The vast majority of enriched pathways are metabolic pathways, particularly those involved in the carbohydrate metabolism, while just few of them are involved on other metabolic processes such as lipid, energy or amino acid metabolism. Pathways involved in genetic information processing were: RNA polymerase, ribosome, and proteosome. These pathways are crucial for transcription, translation and protein folding sorting and degradation, respectively. Finally, the peroxisome was the only pathway enriched involved in cellular processes, particularly in transport and catabolism (Fig 4 D&E). We also observed that the carbohydrate related pathways, along with proteosome, peroxisome and fatty acid degradation were suppressed in conidia coming from high sucrose environment. We also explored the occurrence of alternative splicing events and found 32 cases in total, of which only 17 were in annotated genes. Due to the small number of annotated genes enrichment analysis of alternatively spliced sites was not possible (see supplementary information and supplementary table S6).

#### Epigenetic mechanisms

To explore are the parental effects based on epigenetic processes, we searched for the silver spoon effect using three mutant lines: ∆*dim-2*, which is deficient in DNA methylation; ∆*qde-2*, which is deficient in small RNA processing; and ∆*set-7*, which is deficient in histone 3 lysine 27 trimethylation. All three strains showed the silver spoon effect; initial colony size was bigger when the fungus had experienced 1.5% sucrose in the previous generation (Fig 5; ∆*dim-2* = 1.486 [1.125, 1.805]; ∆*qde-2* = 1.658 [1.401, 1.906]; ∆*set7* = 1.148 [0.743, 1.558]). This suggests that the silver spoon effect is not based on any of these epigenetic mechanisms.

**Figure 5:**
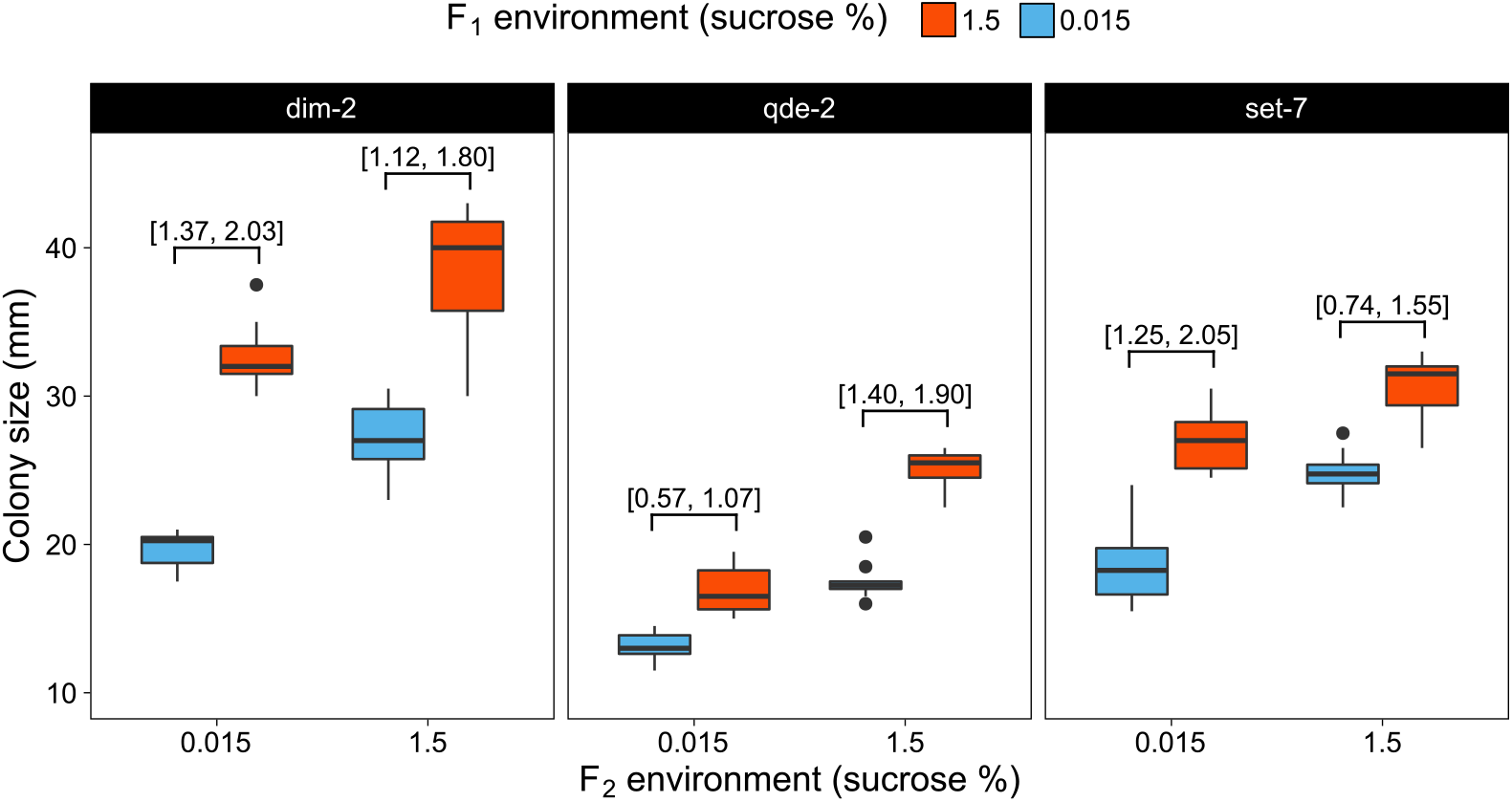
Effect of F_1_ sucrose concentration on deletion mutant strains. Raw data of initial colony size for three mutant strains: ∆*dim-2*, ∆*qde-2* and ∆*set-7*. Numbers inside square brackets are the 95% HPDI of the differences between treatments obtained from model S1 for scaled data.

To further understand the role of epigenetics in the silver spoon effect, we examined in which domains the 379 genes that belonged to the GSEA enriched pathways were located. We observed that all genes belonging to the main GSEA pathways, were located in euchromatic regions that were associated with H3K36me2. Twenty genes in total overlapped with H3K27me3 domains, from these 20 genes, 16 completely overlapped and 4 partially overlapped with H3K27me3 domains. 14 genes overlapped with H3K9me3, of which only one completely overlapped H3K9me3 (Fig 4C). No genes belonging to the enriched pathways exclusively overlapped heterochromatic regions.

## Discussion

Parental effects are a potential mechanism by which organism can deal with environmental challenges [8, 7, 9, 1]. However, our understanding about parental effects still has important limitations. First, it is crucial to investigate how widely distributed parental effects are across taxa, since research so far has mostly focused on animals and plants neglecting other eukaryotes such as fungi. Second, even though parental effects are widely studied their mechanisms are rarely investigated. So far, to our knowledge, there is only one published investigation on parental effects in fungi [20] where maternal investment during sexual cycle of *N. crassa* was explored. Our study is the first one to look into parental effects induced by the environment in fungi, and an in depth investigation of this phenotype, mechanism, and fitness consequences.

Silver spoon effects are those life long fitness advantages that an organism may have because of the access to abundant resources by its parents or during early development [39, 40, 37]. In this case, the fungus in favorable environments had access to more resources that it invested in the next generation (i.e spores). We demonstrated that even if the carry over effect was only relevant during initial growth, it can increase fitness and thus has the potential to be adaptive. Silver spoon effects have been classified by some authors as non-adaptive [41]. However, Bonduriansky and Crean [37] argued that silver spoon effects can indeed enhance parental fitness by increasing the performance of the next generation. Furthermore, Bonduriansky and Crean [37] argued that it is expected that net selection favors silver spoon effects because even though individuals in low conditions will produce low quality descendants and lose fitness, silver spoon effects will naturally increase fitness of high conditions individuals and therefore will enhance fitness on average [37].

Variation in the parental environment can frequently result in some degree of unavoidable transmission of the parental condition, leading to a silver spoon effect. However, apart from the environmental variation, parental investment can vary due to a number of inherent characteristics, such as genetic background, health or age. For example in *Daphnia*, offspring of clonal females that were under the same environmental conditions, considerably differed in life history traits such as, size at birth, age of maturity and number of offspring [42]. To investigate if the silver spoon effect described here is an inevitable consequence of the parental environment, we would need to establish if strains with different genetic backgrounds differ on their efficiency to transfer parental resources. If selection favors increased offspring investment traits, such as storage of metabolic resources and efficiency of cellular processes, these are likely to evolve a variety of strategies for parental investment [37].

To understand the scope that silver spoon effects can have in natural populations, it is crucial to understand their mechanisms. Glycogen serves as a carbon and energy reserve, and glucose as the main energy source in N. crassa [43, 44, 45]. Cultures that were grown under low sucrose conditions were limited by the amount of glucose in the medium, as spore production was severely limited. In addition, the spores produced by a mycelium in 0.015% sucrose had lower glycogen and glucose levels. The glycogen storage and glucose availability in the spores gives a fitness advantage to the fungus, even if in the next generation it grows in a low sucrose environment.

In conjunction with sugar content in the spores, we found that the silver spoon effect involved a dramatic gene expression change, in which pathways related to sugars and carbohydrate metabolism were over-expressed in conidia that experienced 0.015% sucrose. These results are explained by the carbon catabolite repression, a common process among fungi, where the production of enzymes responsible for degrading plant cell wall material is inhibited while preferred carbon sources (e.g sucrose), are available in the environment.

*In nature N. crassa* grows on dead plant material, thus, it heavily relies on breaking down the plant biomass components [46, 47, 48]. For this reason *N. crassa* has a vast enzymatic toolkit that allows it to utilize the variety of simple or complex carbon sources present in the plant cell wall. However, it would be disadvantageous to produce enzymes to break down nutrients that are not available in the substrate [46]. To avoid such costs, *N*.*crassa* has evolved systems to accurately detect the nutrients available in the environment to produce only the needed enzymes [46, 49, 50]. When sucrose is present, the carbon catabolite repression silences the expression of lignocellulolytic genes [46]. When sucrose is not available, the carbon catabolite repression is diminished causing elevated levels of lignocellulolytic genes expression allowing a small secretion of a vast number of different enzymes that allow the fungus to utilize alternative carbon sources [49]. This produces gene expression patterns in which the fungus expresses ribosomal proteins and functional categories related with primary metabolism pathways in sucrose rich environments, while under glucose starvation fungus expresses sugar and carbohydrate metabolism related pathways [51, 48]. This metabolic behavior has been previously observed in *N. crassa* and other fungal species [52]. The mycelium of the next generation will directly germinate from the conidia, therefore the mRNA content of the spores impacts the performance of the next generation.

Similar silver spoon effects were also present when *N. crassa* grew on media containing arabinose, cellulose, or lactose, and absent when it grew on maltose and xylose media. A possible explanation for this might be that the first three environments represent a disadvantage over the sucrose environment and they will trigger the carbon catabolite repression. For example, although cellulose is one of the main plant cell wall components, it is very difficult to degrade, lactose is slowly metabolized [53, 54], and arabinose rewires the fungal cell metabolic pathway triggering a similar response to carbon starvation conditions [55]. On the contrary, maltose and xylose are not very challenging environments, xylose is one of the preferred carbon sources [49] and maltose is actually commonly used as a banding media when studying circadian rhythms [56].

We observed that strains deficient in different epigenetic mechanisms did not prevent silver spoon effects from occurring. DNA methylation and H3K27 trimethylation are associated with heterochromatic regions in *N. crassa*, which have low gene density and expression levels [57, 33]. Most genes belonging to the pathways showing differential expression were associated with euchromatic regions. It appears that carbohydrate metabolism is not under strict epigenetic control in *N. crassa*.

Finally we want to stress the importance of expanding the taxonomic representation on parental effects research and to investigate their adaptive potential even if they are short lived. In comparison to anticipatory effects, silver spoon effects have been widely overlooked even that some of their aspects suggest they might the most widespread type of parental effect across taxa [37]. Contrary to anticipatory effects, silver spoon effects do not depend on the environment predictability, nor on complex mechanisms to asses the environment and adjust the offspring phenotype accordingly. Silver spoon effects may influence the ecology and evolutionary processes in several eukaryotes across the tree of life.

## Acknowledgments

This study was supported by a grant from the Academy of Finland (no. 321584) to IK. We thank the Finnish CSC-IT Center for Science Ltd. for providing computational resources.

## Author contributions

I.K. and M.V. conceived the study. M. V., P.A.M.S., N. N. M., and I.K performed experiments. M.V., I.K., M.V., and P.A.M.S. analyzed the data. M.V and I.K. wrote the manuscript. All authors edited the final manuscript.

## Data access

RNA-sequencing data has been deposited to the short read archive, project number PRJNA907747, with sequence accession numbers SRX18465547–SRX18465558. Other data and scripts are available at https://github.com/mariana19901990/Neurospora-crassa-Parental-effects.

## Supplementary information

Supplemental information for article: Parental effects in a filamentous fungus: phenotype, fitness, and mechanism. By Mariana Villalba de la Peña, Pauliina A. M. Summanen, Neda N. Moghadam, and Ilkka Kronholm.

### Supplementary methods

#### Fitness consequences of parental effects

To investigate fitness consequences of the parental effect we performed a competition experiment. *N. crassa* grew in 1.5 or 0.015%, sucrose concentration for two generations, then 5 000 conidia from each competitor were inoculated to an agar slant with a sucrose concentration of either 1.5 or 0.015%, giving 10 000 conidia in total. When the culture produced conidia, a sample was transferred to a new slant with the same sucrose concentration, and the rest of the conidia were harvested, DNA was extracted, and HRM-PCR was performed to determine the proportion of the marked strain (Fig S4A). See Kronholm et al. for primers, PCR conditions, and DNA extraction from conidia [25]. Two competition experiments were performed, in the first experiment competition was done for 2 transfers and in the second experiment only for 1 transfer. In the first experiment each combination was repeated for 5 times, with 8 combinations and 2 assay environments there were 80 populations. In the second experiment there were 5 replicates per population, 6 combinations, and 2 assay environments giving 60 populations. The combined data contained 140 populations in total.

#### Protein and carbohydrate content in spores

To measure total protein content we used the BCA protein assay kit (ThermoScientific) according to manufacturer instructions. Harvested conidia from F_2_ cultures were counted using CASY cell counter and washed with water to remove any VM medium traces. For protein extraction 40 million conidia were resuspended in 100 µL of lysis buffer (8.2 mL water, 500 µL of 1 M HEPES, 180 µL of 5 M NaCl, 20 µL of 0.5 M EDTA, and 1 mL of 10% Triton-X100) with protease inhibitor (1X). Samples were tranferred to 2 mL tubes containing 0.5 mm diameter glass beads. Using the Omni bead ruptor we lysed the tissue at 0 degrees (3 cycles of 45 second cycles at a speed of 6 m*/*s with a 30 seconds interval). To measure glycogen and glucose content in spores we used the glycogen assay kit (Sigma-Aldrich, MAK016) and the glucose assay kit (Sigma-Aldrich, MAK263) as indicated by the manufacturer. Extraction of glycogen and glucose was performed as described for the protein extraction but 70 million spores were used and they were resuspended in water instead of lysis buffer.

#### RNA-seq and analysis

To investigate the mechanisms behind the parental effects we performed RNA-seq on conidia from the two sucrose condition. After two generations of *N*.*crassa growing in* rich and poor sucrose environmnet, conidia were harvested, filtered and suspended in 5 mL of 0.01% Tween-80. To lyse the tissue we added the cell suspension to 2 mL tubes containing 0.5 mm diameter glass beads and 1 mL of Trizol. Then we processed samples in the Omni bead ruptor for two 30 second cycles at a speed of 6 m*/*s with a 45 second interval. After tissue homogenization, we extracted RNA following Kramer [26]. The purity, concentration and integrity of the extracted RNA was assessed using Nanodrop, Quibit RNA Broad range Kit and the Aligent RNA ScreenTape Analysis (Supplementray table S3). We spiked the total RNA with Ambion ERCC RNA spike-in mixes as specified by the manufacturer [27]. ERCC RNA Spike-In controls consists of two mixes of 92 polyadenylated transcripts with known concentrations. These serve as external controls that facilitate the normalization and performance assessment of RNA-seq data [27]. We added Mix 1 to 1.5% glucose samples and Mix 2 to 0.015% glucose samples. Finally, six biological replicates from each sucrose concentration were sent for to Novogene for mRNA poly A enrichment library preparation, and for transcriptome sequencing using the Illumina NovaSeq platform with 150 bp paired-end libraries.

External controls have shown to be a reliable option to accurately normalize RNA-seq data [27]. However, data normalization that solely relays on ERCC spike-in controls can be risky as they can also be affected by variation coming from library preparation or other sources of unwanted variation [30]. For this reason it is necessary to examine the performance of the spike-in controls. In our data theoretical concentrations of spike-in controls coincide well with the number of transcript counts (Fig S5A). However, the proportion of reads mapping to the ERCC spike-ins were highly variable between libraries (Fig S5B), also in some of the samples, the genes and controls were differently affected by unwanted variation (Fig S6). Based on the control’s performance, we decided to calculate the unwanted variation based on the external spike-in controls. First, we normalized the RNA-seq data using the Trimmed Mean of the M-values approach (Fig S5C), then we calculated the unwanted variation using the RUVg function, from the bioconductor package RUVseq (R environment version 4.0.2), [30]. We used DESEq2 [63] to identify differentially expressed genes and Cluster profiler [58] to perform over representation analysis (ORA) using all annotated genes as universe and gene set enrichment analysis (GSEA), both identifying KEGG pathways. In DE (differential expressed) and enrichment analysis we used Benjamini and Hochberg for multiple testing correction [64].

#### Alternative splicing detection

Besides gene expression we also looked for the existence of different alternative splicing events between treatments. We ran rMATS [65] considering each sucrose environment as a treatment and each sample as a replicate. We specified a read length of 150 bp and the type of reads as paired. Using the bioconductor package maser [66] we filtered the rMATS junction count output (strict output as it only counts the junction reads) to obtain only those events that were cover with a minimum of 20 reads, false discovery rate smaller than 0.01 and a percent.spliced-in (PSI) difference of at least 0.2. The PSI index indicates the ratio between reads including or excluding sequences of interest (e.g exons; [67]). A PSI equal to 1 indicates sequences that are included in all transcripts. PSI values below 1 imply reduced inclusion of alternative sequences and indicated the percentage of proteins that contain the sequence compared to the total transcript population [68, 67].

We found 32 events of alternative splicing. Three of them were alternative 3’ splice sites, three alternative 5’ splice sites, only one event was a skipped exon and 25 were retained intron events. We did not find enough alternative splicing events to do further enrichment analysis. From the 32 spliced genes 17 were annotated, the rest were described as hypothetical proteins. The main function of the annotated proteins were mainly related with, kinase activity, transcription regulation and cell structure (Table S6)

#### Epigenetic mechanisms

We wanted to explore if the GSEA gene set are related to the genome domains affected by the mutant strains ∆*dim-2* and ∆*set-7. dim-2* encodes a methyltrasnferase responsible of DNA methylation, which in turn is associated with H3K9me3 domain. *set-7* regulates the H3K27m3. H3K36me2 is an opposing domains as it usually don not overlap with K3K27me3 and H3K9me3. We first obtained ChIP-seq reads for H3K9me3 (accession number SRX248101) and H3K27me3 (accession number SRX248097) from Jamieson et al. (2013) [33], and H3K36me2 (accession number SRX4549854) from Bicocca et al. (2018) [70]. Reads were aligned to the reference genome using BWA, and duplicate reads were removed by Picard tools. Domains of histone modifications were identified using RSEG 0.4.9 [71]. Using bedtools we identified the intersecting regions between each histone modification domains and the genes from each GSEA enriched pathway. We considered a gene to belong to a histone modification domain if at least 20% of the gene overlapped with the histone modification domain.

#### Statistical analysis of existence and duration of parental effects

The model for analyzing existence and duration of parental effects was:

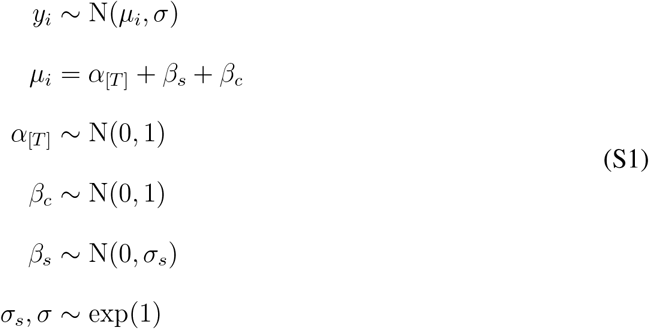

where *y*_*i*_ is *i*th observation of initial colony size, *α*_[*T*]_ is the intercept for each treatment, *β*_*c*_ conidial viability (number of colonies in sorbose plates), and *β*_*s*_ the slant effect. The treatment summarized parental and current environmental conditions as specified in table S2. Similar model was also used to examine the effect of treatment on spore size, viability, protein and sugar content in which case *y*_*i*_ was spore diameter, number of colonies, protein, glycogen and glucose amount, respectively. For MCMC estimation four independent chains were run, with 1 000 warm-up iterations, followed by 4 000 samples. We ran the models using specific informative *α, β* ∼ N(0, 1) and weakly informative priors *α, β ∼* N(0, 5). However, both priors resulted in the same model output. The traceplots showed that the model converged and no divergent transitions were found, 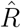 values were never higher than one.

#### Statistical analysis of fitness consequences of parental effects

The final model used to estimate fitness effects from competition experiments was:

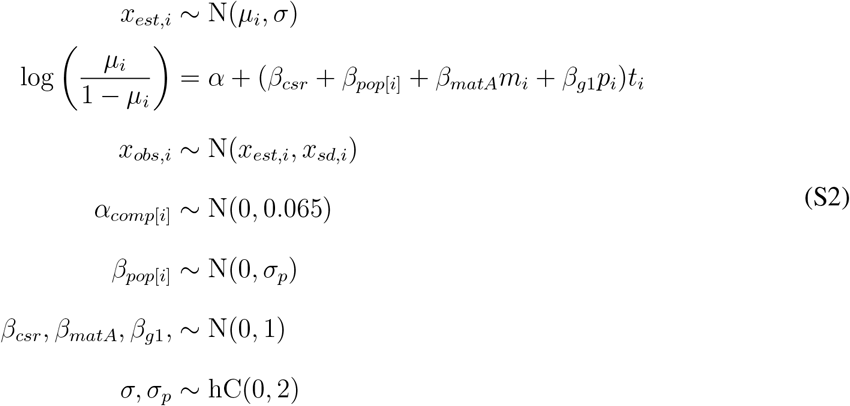

where *x*_*obs,i*_ is the *i*th observed marked strain proportion, *x*_*sd,i*_ is the *i*th observed uncertainty for that observation, *x*_*est,i*_ is the *i*th estimated proportion, *α* is the intercept, *β*_*pop*[*i*]_ is the slope effect for each population, *β*_*csr*_ is the effect of the *csr-1\** allele, *β*_*matA*_ is the effect of mating type A, *m*_*i*_ is an indicator whether the marked strain is *mat A, β*_*g*1_ is the effect of the parental environment, *p*_*i*_ is an indicator about the parental environment of the marked strain, *t*_*i*_ is the transfer number, *σ*_*p*_ is standard deviation among populations, and *σ* is the error standard deviation. The indicator for mating type, *m*_*i*_ *∈* {*−*1, 1}, gets a value of 1 when the marked strain is *mat A*, and *−*1 when the marked strain is *mat a*. The indicator for parental environment, *p*_*i*_ *∈* {*−*1, 0, 1}, gets a value of 1 when parent of the marked strain comes from 1.5% sucrose and unmarked strain from 0.015%, value of *−*1 when the situation is reversed, and 0 when parents of both strains grew in the same environment. We used weakly regularizing priors for slope effects, and an informative prior for the intercept, since all competitions were started with a frequency of 0.5 of the marked strain. MCMC estimation was done using two chains, with 1 000 warmup iterations and then 4 000 sampling iterations. The model converged: all parameters had 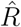 values of 1, trace plots showed that all chains converged to the same solution, and no problems with divergent transitions were encountered. Since slope effects represent the log relative fitness in this model, posterior distributions of slope effects were transformed to relative fitness by expression *W* = exp(*β*).

## Supplementary tables

**Table S1:** Sample size and biological replicates. Shown are the sample size (n) and the number of biological replicates used in each treatment. The biological replicates refers to the number of slants used in each treatment.

| Experiment | Generation | n | Biological replicates |
| --- | --- | --- | --- |
| Experiment 1 | F2 | 40 | 3 |
| Experiment 2 | F2 | 40 | 3 |
| Experiment 3 | F2 | 40 | 3 |
| Experiment 4 | F2 | 40 | 3 |
| Experiment 5 | F2 | 40 | 3 |
| Experiment 6 | F2 | 60 | 5 |
| Experiment 7 | F2 | 36 | 9 |
| Experiment 8 | F2 | 26 | 6 |
| Experiment 9 | F2 | 28 | 7 |
| Experiment 1 | F3 | 40 | 3 |
| Experiment 2 | F3 | 40 | 3 |
| Experiment 3 | F3 | 40 | 3 |
| Experiment 5 | F3 | 40 | 5 |
| Experiment 6 | F3 | 60 | 5 |

**Table S2:** Condensation of the sucrose environments into treatments. Here we show the summarized sucrose environments into four treatments. Such treatments where used as a predictor in the model specified in S1.

| Treatment | F<sub>1</sub> sucrose environment | F<sub>2</sub> sucrose environment | F<sub>3</sub> sucrose environment |
| --- | --- | --- | --- |
| Treatment 1 | 1.5% | 1.5% | 1.5% |
| Treatment 2 | 1.5% | 0.015% | 0.015% |
| Treatment 3 | 0.015% | 0.015% | 0.015% |
| Treatment 4 | 0.015% | 1.5% | 1.5% |

**Table S3:** Summary of the sequenced RNA samples quality and alignment metrics.

| Sample ID | A 260/280 | A 260/230 | RIN | Number of reads | Mapped reads |
| --- | --- | --- | --- | --- | --- |
| S1 1.5% | 2.11 | 2.13 | 7.7 | 13419881 | 96.70% |
| S5 1.5% | 2.05 | 1.25 | 7.4 | 13588548 | 95.75% |
| S8 1.5% | 2.09 | 1.88 | 7.5 | 14622588 | 95.63% |
| S7 1.5% | 2.09 | 1.81 | 7.5 | 15453849 | 95.54% |
| S2 1.5% | 2.09 | 2.01 | 7.5 | 14922835 | 96.04% |
| S9 1.5% | 2.10 | 1.93 | 7.7 | 12557867 | 95.50% |
| S8 0.015% | 2.09 | 1.99 | 9.0 | 12640968 | 93.86% |
| S4 0.015% | 2.09 | 2.01 | 8.7 | 11205530 | 93.75% |
| S5 0.015% | 2.10 | 2 | 9.0 | 13457509 | 94.34% |
| S6 0.015% | 2.09 | 1.81 | 9.0 | 11575198 | 94.51% |
| S7 0.015% | 2.08 | 2.09 | 9.0 | 14056380 | 93.86% |
| S9 0.015% | 2.10 | 2.10 | 9.0 | 12468794 | 94.20% |

**Table S4:** Experimental design of competition experiment. Strains were competed in different combinations to estimate independent effects for the marker, mating type, and parental environment. The strains were developed previously in [25], and they are nearly isogenic. Fungal Genetics Stock Center IDs are: 2489 *mat A* = B 26708, 2489 *mat a* = B 26709, 2489 *mat A csr-1\** = B 26710, 2489 *mat a csr-1* = B 26711.

| Strain 1 | Strain 1 parental env | Strain 2 | Strain 2 parental env | Competition env |
| --- | --- | --- | --- | --- |
| 2489 *csr-1\** *mat A* | 1.5% | 2489 *mat a* | 1.5% | 1.5% |
| 2489 *csr-1\** *mat A* | 1.5% | 2489 *mat a* | 0.015% | 1.5% |
| 2489 *csr-1\** *mat a* | 1.5% | 2489 *mat A* | 0.015% | 1.5% |
| 2489 *mat A* | 1.5% | 2489 *csr-1\** *mat a* | 0.015% | 1.5% |
| 2489 *mat a* | 1.5% | 2489 *csr-1\** *mat A* | 0.015% | 1.5% |
| 2489 *csr-1\** *mat a* | 1.5% | 2489 *mat A* | 1.5% | 1.5% |
| 2489 *csr-1\** *mat A* | 0.015% | 2489 *mat a* | 0.015% | 0.015% |
| 2489 *csr-1\** *mat A* | 1.5% | 2489 *mat a* | 0.015% | 0.015% |
| 2489 *csr-1\** *mat a* | 1.5% | 2489 *mat A* | 0.015% | 0.015% |
| 2489 *mat A* | 1.5% | 2489 *csr-1\** *mat a* | 0.015% | 0.015% |
| 2489 *mat a* | 1.5% | 2489 *csr-1\** *mat A* | 0.015% | 0.015% |
| 2489 *csr-1\** *mat a* | 0.015% | 2489 *mat A* | 0.015% | 0.015% |

**Table S5:** Results of model described in equation S1. The models analyze the effect of F_1_ sucrose concentration on initial growth, spore size, viability on generation two and three in the wild type and mutant strains.*α* is the intercept for each treatment, *β*_*c*_ conidia viability and *β*_*s*_ controlling for the slant effect. Only the estimates of the fixed effects are reported.

| Response variable | Model terms | Estimate [95% HPDI] |  |  |  |  |
| --- | --- | --- | --- | --- | --- | --- |
| | | $\alpha_{T1}$ | $\alpha_{T2}$ | $\alpha_{T3}$ | $\alpha_{T4}$ | $\beta_c$ |
| $F_2$ initial growth | $\alpha_T + \beta_s$ | 0.985 [0.799, 1.173] | -0.01 [-0.196, 0.178] | -0.728 [-0.919, -0.547] | -0.018 [-0.376, -0.005] | NA |
| $F_2$ initial growth | $\alpha_T + \beta_s + \beta_c$ | 0.916 [0.621, 1.189] | -0.087 [-0.362, 0.199] | -0.600 [-0.885, -0.317] | -0.209 [-0.473, -0.086] | 0.034 [-0.149, 0.222] |
| $F_3$ initial growth | $\alpha + \beta_s + \beta_c$ | 0.672 [0.490, 0.871] | -0.730 [-0.926, -0.544] | -0.490 [-0.688, -0.308] | 0.550 [0.352, 0.732] | 0.188 [0.071, 0.291] |
| $F_2$ spore size | $\alpha_T$ | 0.009 [-0.239, 0.249] | -0.009 [-0.241, 0.244] | NA | NA | NA |
| $F_2$ viability | $\alpha_T + \beta_s$ | -0.051 [-0.438, 0.322] | 0.122 [-0.288, 0.530] | NA | NA | NA |
| $F_3$ spore size | $\alpha_T$ | -0.118 [-0.580, 0.347] | -0.218 [-0.677, 0.232] | 0.032 [-0.437, 0.488] | 0.304 [-0.154, 0.783] | NA |
| $F_3$ viability | $\alpha_T + \beta_s$ | 0.548 [0.089, 0.959] | 0.370 [-0.807, 0.056] | -0.128 [-0.542, 0.279] | 0.357 [-0.052, -0.766] | NA |
| $\Delta dim-2$ initial growth | $\alpha_T$ | 1.176 [0.647, 1.748] | 0.419 [-0.220, 0.868] | -1.287 [-1.780, -0.712] | -0.307 [-0.722, -0.166] | NA |
| $\Delta qde-2$ initial growth | $\alpha_T$ | 1.509 [1.325, 1.685] | -0.267 [-0.449, -0.087] | -1.091 [-1.278, 0.913] | -0.149 [-0.325, 0.030] | NA |
| $\Delta set-7$ initial growth | $\alpha_T$ | 1.070 [0.775, 1.361] | 0.336 [0.029, 0.613] | -1.331 [-1.617, -1.031] | -0.079 [-0.361, 0.201] | NA |
| cellulose initial growth | $\alpha_T$ | 1.927 [1.366, 2.416] | 1.496 [0.973, 2.056] | NA | NA | NA |
| arabinose initial growth | $\alpha_T$ | 1.439 [0.805, 2.064] | 1.841 [1.198, 2.462] | NA | NA | NA |
| lactose initial growth | $\alpha_T$ | 1.211 [0.567, 1.877] | 1.809 [1.159, 2.451] | NA | NA | NA |
| maltose initial growth | $\alpha_T$ | -0.871 [-1.994, 0.269] | -0.589 [-1.831, 0.652] | NA | NA | NA |
| xylose initial growth | $\alpha_T$ | -0.414 [-1.496, 0.777] | -0.027 [-1.146, 1.138] | NA | NA | NA |
| time 1 growth rate | $\alpha_T + \beta_s$ | 1.248 [0.854, 1.665] | 0.630 [0.216, 1.017] | NA | NA | NA |
| time 2 growth rate | $\alpha_T + \beta_s$ | 0.099 [0.248, 0.459] | 0.312 [-0.048, 0.670] | NA | NA | NA |
| time 3 growth rate | $\alpha_T + \beta_s$ | -0.018 [-0.409, 0.334] | 0.083 [-0.269, 0.453] | NA | NA | NA |
| number of conidia | $\alpha_T$ | 1.743 [1.519, 1.974] | NA | NA | NA | NA |
| protein content | $\alpha_T$ | -0.10 [-0.39, 0.19] | 0.10 [-0.19, 0.39] | NA | NA | NA |
| glycogen content | $\alpha_T$ | 0.80 [0.45, 1.13] | -0.80 [-1.13, -0.46] | NA | NA | NA |
| glucose content | $\alpha_T$ | 0.90 [0.70, 1.09] | -0.90 [-1.10, -0.70] | NA | NA | NA |

**Table S6:** Alternative splicing events. Shown are the 32 significant alternative slicing (AS) events detected with rMATS. The AS events are categorized as intron retention (RI), skipped exon (SE),alternative 3’ splice sites (A3SS) and alternative 5’ splice sites (A5SS). The percent spliced in (PSI) indicates the efficiency of splicing a specific exon into the transcript population of a gene

| Gene | pValue | FDR | PSI difference | AS event | Gene description | Biological process /molecular function |
| --- | --- | --- | --- | --- | --- | --- |
| NCU09368 | 0 | 0 | 0.316 | RI | cation diffusion facilitator 10 | cellular transition metal ion homeostasis |
| NCU01166 | 0 | 0 | 0.607 | RI | camp-dependent protein kinase regulatory chain | negative regulation of cAMP-dependent protein kinase activity |
| NCU05564 | 0 | 0 | 0.350 | RI | peroxisomal membrane protein PEX31 | peroxisome organization |
| NCU04272 | 0 | 0 | 0.345 | RI | ZZ type zinc finger domain-containing protein, variant | Zinc ion binding |
| NCU01500 | 0 | 0 | 0.457 | RI | nicotinamide riboside kinase 1 | NAD biosynthesis via nicotinamide riboside salvage pathway |
| NCU03855 | 0 | 0 | 0.357 | RI | CCR4-NOT transcription complex | regulation of transcription, DNA-templated |
| NCU00289 | 0 | 0 | 0.454 | RI | TAH-1 | transcription, DNA-templated |
| NCU06110 | 0 | 0 | 0.399 | RI | thiazole biosynthetic enzyme, variant 3 | thiamine biosynthetic process |
| NCU03954 | 1.297e-09 | 8.730e-09 | 0.300 | RI | tbulin gamma chain | mitotic cell cycle |
| NCU07347 | 1.625e-03 | 4.716e-03 | -0.213 | RI | endo-beta-1,3-glucanase | carbohydrate metabolic process |
| NCU05791 | 2.889e-13 | 2.759e-12 | 0.254 | RI | SOM1 protein | positive regulation of transcription by RNA polymerase II |
| NCU06716 | 4.143e-07 | 2.229e-06 | 0.316 | RI | short chain dehydrogenase/reductase, variant | oxidoreductase activity |
| NCU08727 | 0 | 0 | 0.215 | RI | hypothetical protein |  |
| NCU03636 | 0 | 0 | 0.237 | RI | hypothetical protein |  |
| NCU01208 | 0 | 0 | 0.203 | RI | hypothetical protein |  |
| NCU00550 | 1.250e-06 | 6.274e-06 | 0.298 | RI | hypothetical protein |  |
| NCU00700 | 1.443e-15 | 1.857e-14 | 0.310 | RI | hypothetical protein |  |
| NCU03848 | 1.538e-08 | 9.486e-08 | 0.396 | RI | hypothetical protein |  |
| NCU01983 | 1.675e-12 | 1.377e-11 | 0.215 | RI | hypothetical protein |  |
| NCU09994 | 1.709e-14 | 1.874e-13 | 0.217 | RI | hypothetical protein |  |
| NCU05286 | 2.059e-04 | 6.773e-04 | 0.238 | RI | hypothetical protein |  |
| NCU08638 | 2.855e-07 | 1.565e-06 | 0.238 | RI | hypothetical protein |  |
| NCU03882 | 3.685e-14 | 3.636e-13 | 0.363 | RI | hypothetical protein |  |
| NCU01145 | 6.231e-07 | 3.293e-06 | 0.228 | RI | hypothetical protein |  |
| NCU04224 | 8.883e-12 | 6.919e-11 | 0.271 | RI | hypothetical protein |  |
| NCU09995 | 0 | 0 | -0.224 | SE | hypothetical protein |  |
| NCU03804 | 2.083e-13 | 5.209e-12 | 0.232 | A3SS | serine/threonine-protein phosphatase 2B catalytic subunit | Fungal-type cell wall organization |
| NCU00468 | 0 | 0 | 0.397 | A3SS | prephenate dehydrogenase | tyrosine biosynthesis |
| NCU04164 | 0 | 0 | -0.235 | A3SS | hypothetical protein |  |
| NCU10853 | 0 | 0 | 0.309 | A5SS | serine/threonine protein kinase-57 | protein phosphorylation, mRNA cis splicing |
| NCU05791 | 3.819e-10 | 7.202e-09 | 0.238 | A5SS | SOM1 protein | positive regulation of transcription by RNA polymerase II |
| NCU03836 | 1.520e-05 | 1.254e-04 | -0.263 | A5SS | tRNA N6-adenosine threonylcarbamoyltransferase | positive regulation of transcription by RNA polymerase II |

## Supplementary figures

**Figure S1:**
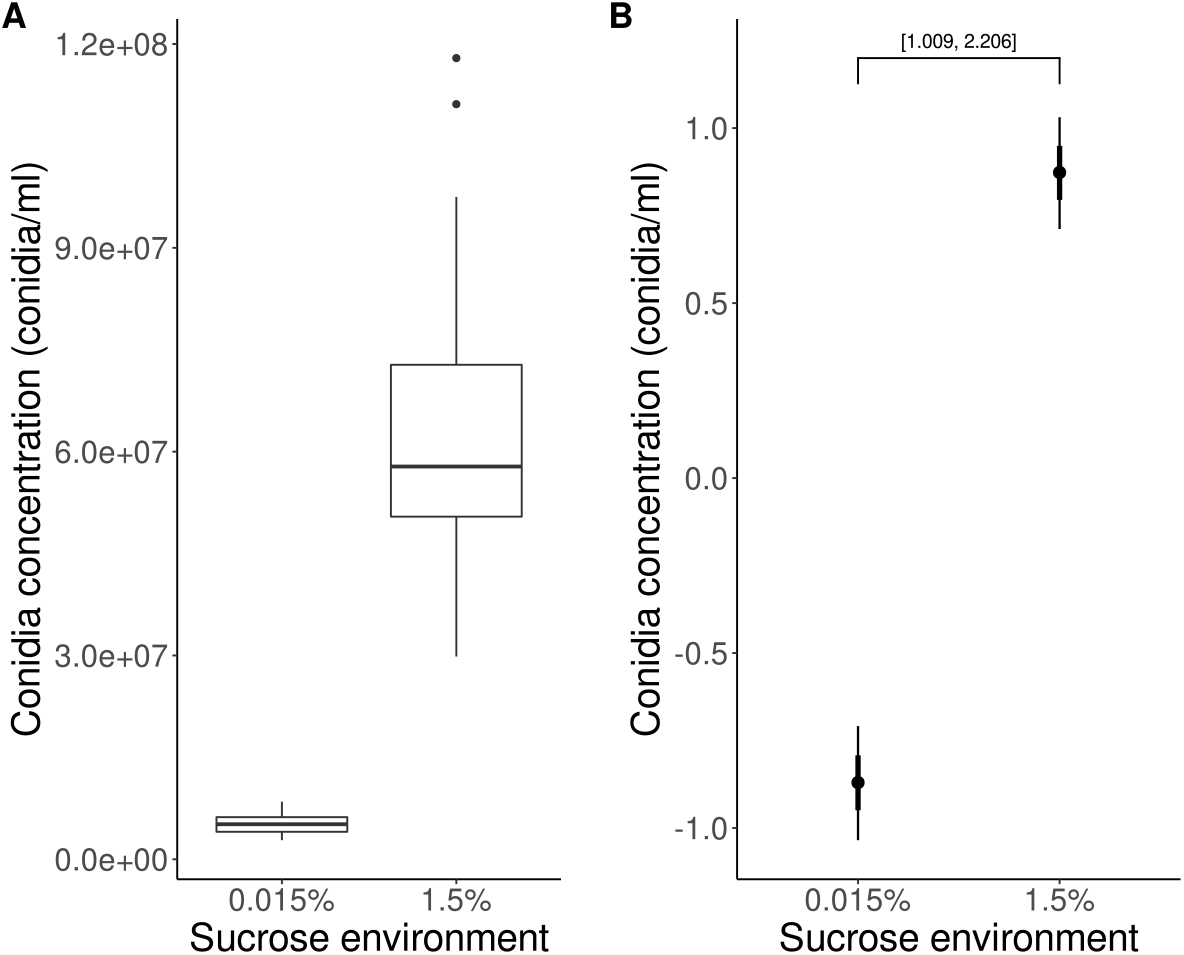
Number of conidia produced in each sucrose concentration. Data from F_2_ and F_3_ samples is combined. **(A)** Raw data. **(B)** Model estimates, the numbers in square brackets represents the 95% HPDI of difference between treatments.

**Figure S2:**
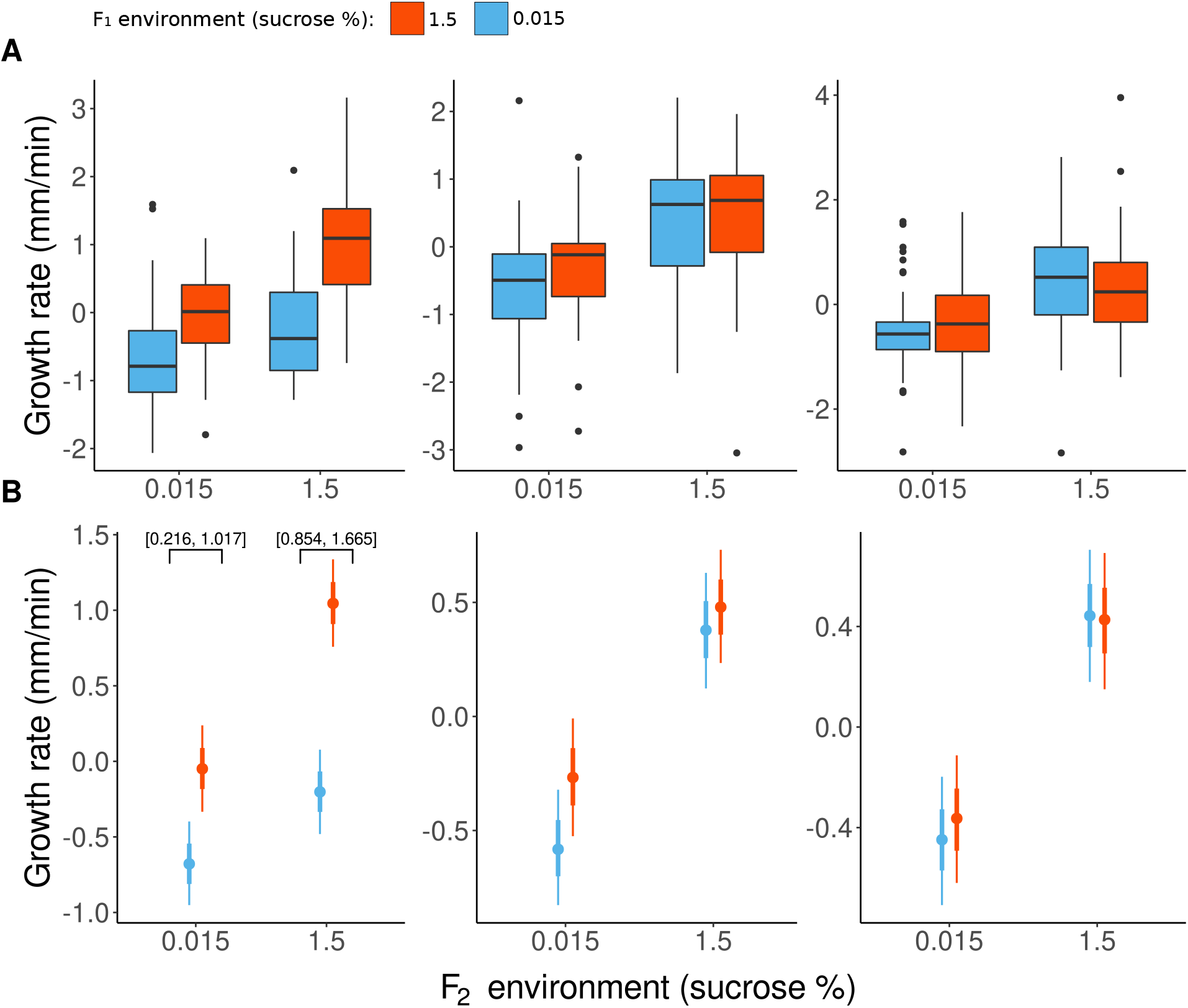
Growth rate of the mycelial colony measured at three time points. First time point (left) was measured 14 to 18 hours after inoculation. Second time point (center) was measured 16-22 after inoculation and the third time point (right) was measured 18-26 hours after inoculation. **(A)** Centered data. **(B)** Model estimates, numbers inside square brackets represent the 95% HPDI of the difference between treatments

**Figure S3:**
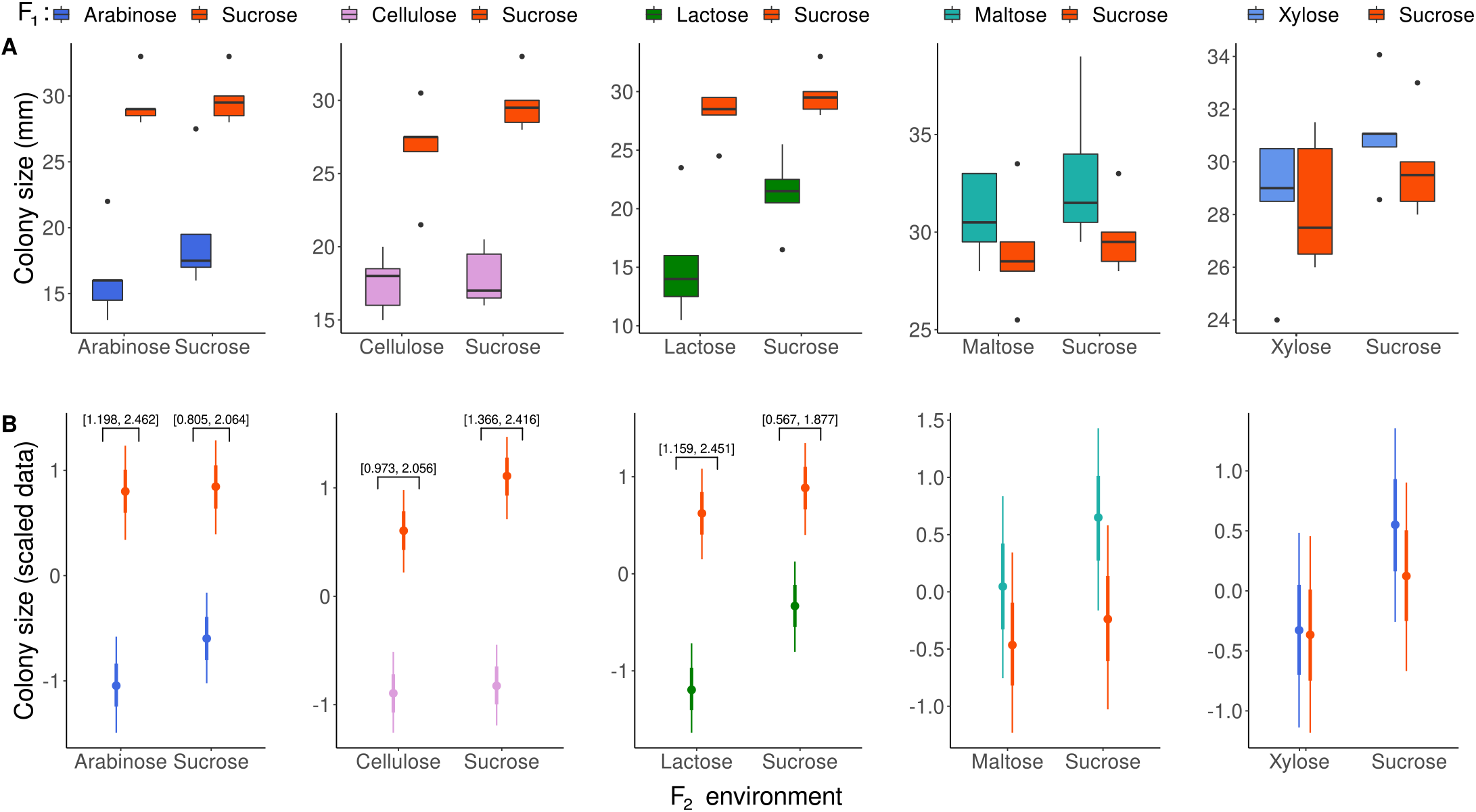
Effect of F_1_ carbon source on F_2_ growth. **(A)** Raw data showing the results of the match-mismatch experiment using various carbon sources. **(B)** Model estimates of initial colony sizes. The numbers in square brackets are the 95% HPDI of differences between treatments.

**Figure S4:**
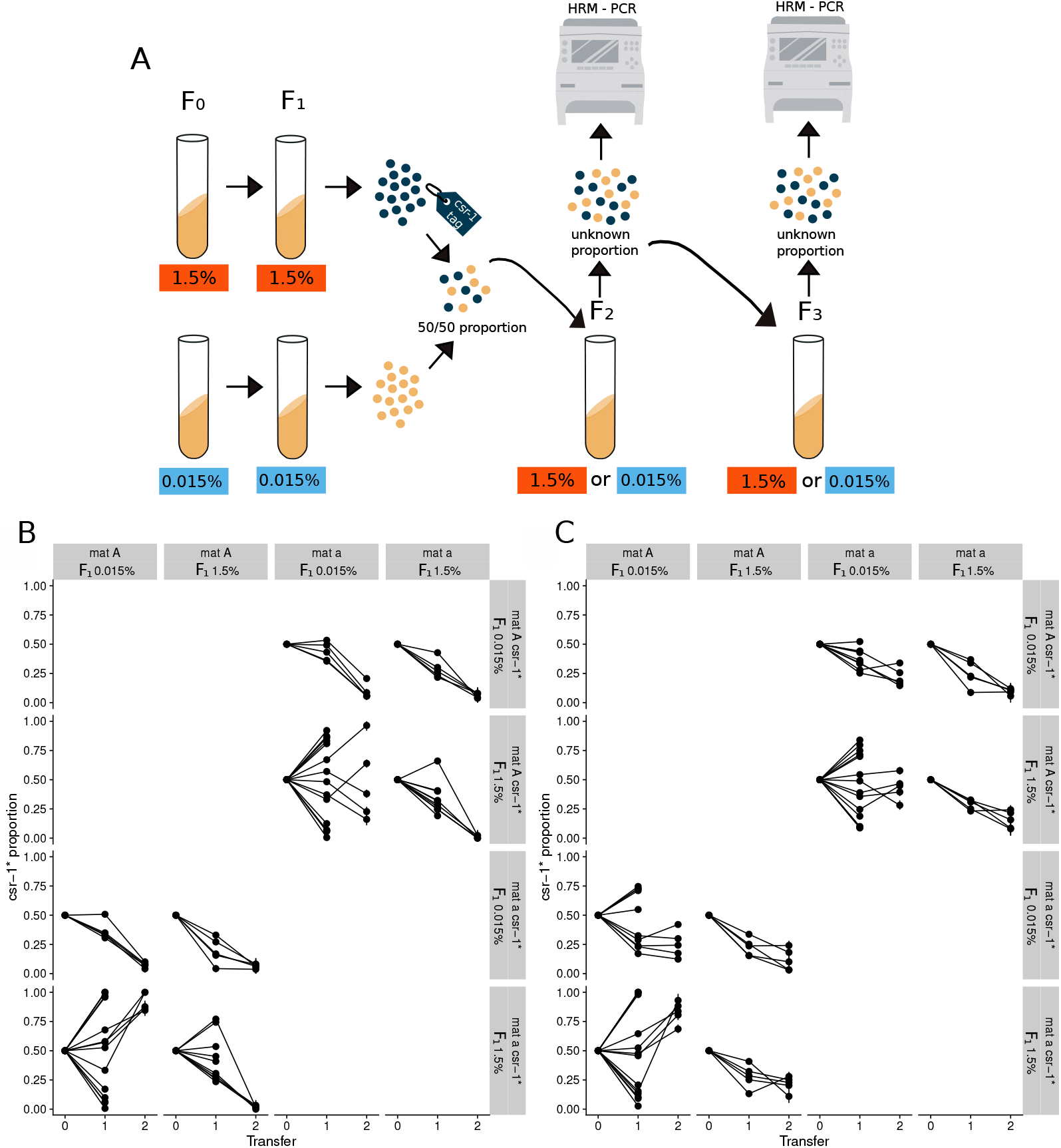
Experimental design and frequency trajectories of the marked strain in competition experiments. **(A)** Diagram of the competition experiment. *N. crassa* grew in slants with 1.5% or 0.015% sucrose. After two generations spores from the two sucrose concentrations were harvested and joined in a single slant and let them compete in both sucrose concentrations. To identify spores coming from each environment, strains with csr-1 tag were used (blue spores). This allowed to determine the proportion of spores produced by each environment strain using HRM-PCR. The experiment was performed for two generations (i.e transfers). **(B & C)** To account for the fitness effect of the csr-tag and the mating type, several competition experiments were performed, in which the csr-1 tag and mating type were combined in eight different ways. Facet labels show strain genotypes and the parental F_1_ environments experienced by the strain (sucrose %). Note that some panels are empty because strains with the same mating type cannot be competed against one another. **(B)** Competitions done in 1.5% sucrose environment. **(C)** Competitions done in 0.015% sucrose environment.

**Figure S5:**
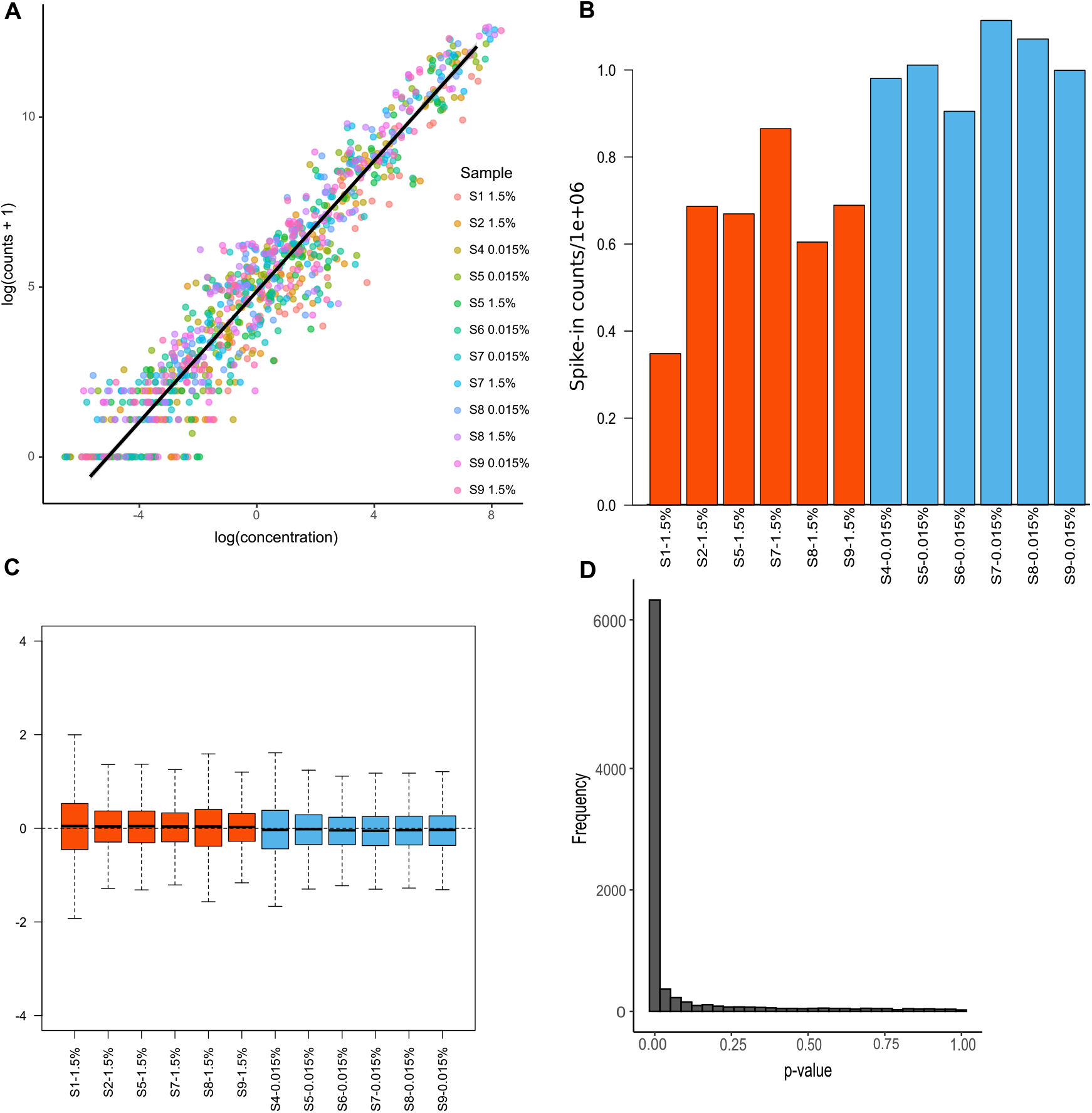
Normalization and validation of RNA-seq data. **(A)** Linear regression of the nominal concentration against the counts obtained of the spike-in controls in all the samples. **(B)** Number of spike-in sequences in each library. **(C)** RLE (relative log expression) graph showing TMM normalization data after removing unwanted variation. **(D)** DESEq2 p-value distribution. Samples coming from 1.5% sucrose environment are presented in orange and samples coming from 0.015% sucrose environment are presented in blue.

**Figure S6:**
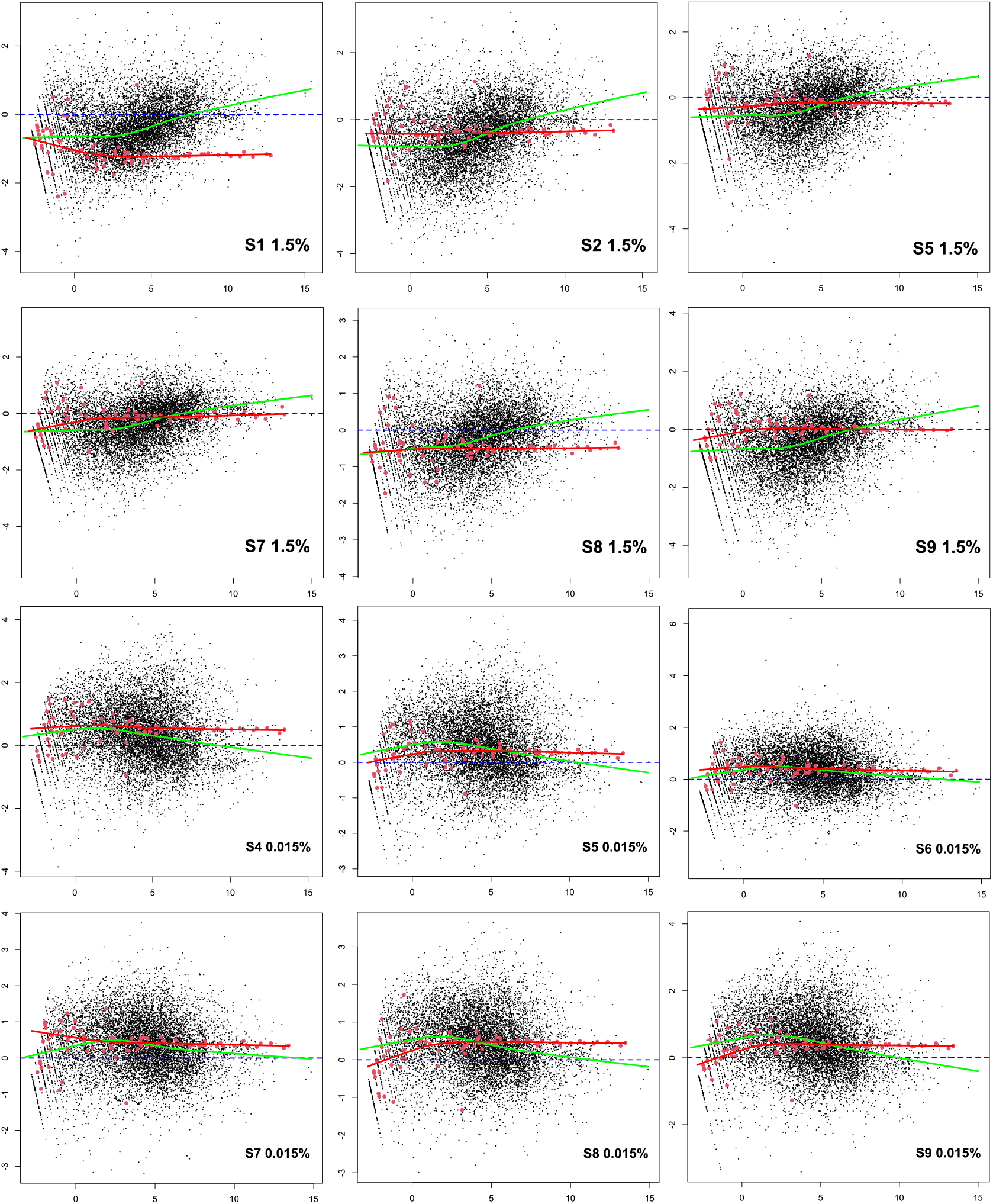
MD graphs. MD plots of unnormalized data. The red points represent the spike-in controls. The red and the green lines represent the output cyclic loess regresion of the spike-in and the genes respectively.

